# The X chromosome of insects predates the origin of Class Insecta

**DOI:** 10.1101/2023.04.19.537501

**Authors:** Melissa A. Toups, Beatriz Vicoso

**Affiliations:** Department of Life and Environmental Sciences, Bournemouth University, Poole BH12 5BB, United Kingdom; Institute of Science and Technology Austria, 3400 Klosterneuburg, Austria

## Abstract

Sex chromosomes have evolved independently multiple times, but why some are conserved for more than 100 million years whereas others turnover rapidly remains an open question. Here, we examine the homology of sex chromosomes across nine orders of insects, plus the outgroup springtails. We find that the X chromosome is shared among all insect orders and springtails; the only exception is in the Lepidoptera, which has lost the X and now has a ZZ/ZW sex chromosome system. Therefore, the ancestral insect X chromosome has persisted for more than 450 million years – the oldest known sex chromosome to date. Further, we suggest that the shrinking of gene content the Dipteran X chromosome has allowed for a burst of sex-chromosome turnover that is absent from other speciose insect orders.

## Introduction

Although sexual reproduction was already present in the ancestor of all animals, the specific mechanisms controlling the development of males and females differs widely between clades. Often, sex chromosomes such as the X and Y of mammals carry the master sex determination genes (Bachtrog et al. 2014). Despite sharing broad characteristics (e.g. the Y is typically gene poor and heterochromatic), these chromosomes are homologs that evolved from autosomal chromosomes independently multiple times across animals, and the processes shaping their convergent differentiation have been studied extensively. While young and undifferentiated sex chromosomes often undergo turnover (i.e. they revert to autosomes, as another chromosome pair takes on the role of sex determination), fully differentiated sex chromosomes are thought to be extremely stable (Pokorná and Kratochvíl 2009). This view of differentiated sex chromosomes as an “evolutionary trap” was largely shaped by model organisms such as eutherian mammals (Pokorná and Kratochvíl 2009; Vicoso 2019), which have all shared the same sex chromosomes for at least 166 million years (Veyrunes et al. 2008). What drives some sex chromosomes to turnover, while others differentiate fully and are maintained over long periods is still unclear. How long sex chromosomes can be conserved for, and what eventually leads to their loss, are also still open questions.

Insects are the most speciose and evolutionary successful animals on the planet, and have colonized almost all terrestrial and freshwater ecosystems (Misof et al. 2014; Stork 2018). The origins of insects is dated as far back as 441 MYA (Misof et al. 2014), either concurring with or shortly after the evolution of land plants (Misof et al. 2014; Morris et al. 2018). Insects use a great diversity of genetic sex-determining mechanisms, including the more evolutionary common systems of XY and ZW sex chromosomes, as well as more rare genetic sex-determining systems such as a paternal genome elimination (PGE) and haplodiploidy (Blackmon et al. 2017). Many orders also have a preponderance of X0 systems, which are relatively rare in other groups of organisms (Bachtrog et al. 2014), and may have been the ancestral state of the clade (Blackmon et al. 2017). Alternatively, these X0 systems may be the result of independent loss of the male-specific Y, which can occur in ancient sex chromosomes (Bachtrog et al. 2014).

Whether this diversity in sex determining systems reflects the independent gain and loss of sex-linked chromosomes, or instead the conservation (and modification) of an ancestral pair of chromosomes, similarly to the mammalian system, has been a longstanding question. An early comparison of *Drosophila melanogaster* (Diptera), *Bombyx mori* (Lepidoptera), and *Tribolium castaneum* (Coleoptera) came to the conclusion that different chromosomal elements have been coopted in these different groups (Pease and Hahn 2012). However, the systematic sequencing of species of various insect orders has shifted this view. First, studies that compared multiple species within orders have detected conservation of the X-linked gene content in Hemiptera (Pal and Vicoso 2015; Mathers et al. 2021), Orthoptera (Xinghua et al. 2022), and Coleoptera (Bracewell et al. 2023), despite the diversity of sex determination systems found in these groups (including X0, XY, multiple X chromosomes, but also PGE and haplodiploidy (Blackmon et al. 2017)). The only currently known exception to the conservation of an X chromosome within an insect order is Diptera, where sex-chromosome turnover from the ancestral element F has been documented extensively involving many different chromosomal elements of the genome (Vicoso and Bachtrog 2015).

Recent studies comparing the gene content of different orders have also suggested that conservation of the X-chromosome may occur over longer periods of time. Homology with the ancestral X chromosome of Diptera (element F) has been detected in Blattodea (Meisel et al. 2019), Odonata (Chauhan et al. 2021), Coleoptera (Chauhan et al. 2021; Xinghua et al. 2022), Hemiptera (Xinghua et al. 2022), and Orthoptera (Xinghua et al. 2022), raising the possibility that the X chromosome has been conserved since the split of Paleopterans and Neopterans 365 million years ago. However, given the sparse distribution of data across the insect phylogeny, and the small number of chromosomes in some clades, it has been unclear if this is simply recruitment of shared gene content in different orders or ancestral homology (Chauhan et al. 2021; Xinghua et al. 2022). For instance, in the best studied insect order, Diptera, conservation of the X chromosome was originally assumed, when the same chromosomal element (called “Muller element A”) was found to correspond to the X of the fruit fly *D. melanogaster* and the mosquito *A. gambiae*, the first two species to be sequenced (Zdobnov et al. 2002). The sampling of more clades showed that a different chromosomal element (“element F”) was ancestrally the X, and that element A was secondarily acquired as the X independently by both fruit flies and mosquitoes (Vicoso and Bachtrog 2015).

Here, we analyze published genome assemblies from nine orders of insects, and the outgroup Collembola, to determine if the X chromosome originated once or multiple times independently. We find that the X chromosome is homologous across eight orders of insects and the outgroup, Collembola. The only exception to this pattern, the Lepidopteran Z chromosome, is not homologous to the X chromosome shared across insects, and the transition to female heterogametey was accompanied by a sex-chromosome turnover. Our results support homology of the X chromosomes across multiple insect orders as the result of ancestry and not independent recruitment of shared genes. Further, we discuss the evolution of shared gene content of the X chromosome in different orders of insect across the phylogeny, and propose that the shrinking of gene content in the Dipteran X chromosome may have led to its turnover, giving rise to the tremendous diversity of XY systems in this group.

## Methods

### Acquisition of genome assemblies and annotations

All species and their sex-chromosome systems used here are listed in Table 1. We inferred the sex-chromosome systems for each species using the Tree of Sex database (The Tree of Sex Consortium et al. 2014).

**Table 1.** Species analyzed in this study and their sex-chromosome system.

| Order | Species | Common name | Male Karyotype |
| --- | --- | --- | --- |
| Plecoptera | *Isoperla grammatica* | common yellow sally | 26(24A+X<sub>1</sub>X<sub>2</sub>) |
| Orthoptera | *Schistocerca americana* | American grasshopper | 23(22A+X) |
| Phasmatodea | *Timema cristinae* | Cristina's Timema | 25(24A+X) |
| Lepidoptera | *Bombyx mori* | silkworm | 56(54A+ZZ) |
| Diptera | *Lucilia cuprina* | Australian sheep blowfly | 12(10A+XY) |
| Coleoptera | *Coccinella septempunctata* | seven-spotted ladybird | 20(18+XY) |
| Neuroptera | *Chrysoperla carnea* | common green lacewing | 14(12+XY) |
| Hemiptera | *Acyrtosiphon pisum* | pea aphid | 7(6A+X) |
| Odonata | *Ischnura elegans* | blue-tailed damselfly | 27(13A+X) |
| Collembola | *Sinella curviseta* | slender springtail | 11(5A+X) |
| Collembola | *Allacma fusca* | globular springtail | 10(4A+X<sub>1</sub>X<sub>2</sub>) |

Chromosome-level genome assemblies and annotations were downloaded from the National Center for Biotechnology Information (NCBI) website for seven orders of class Insecta, including the blue-tailed damselfly (Insecta: Odonata), *Ischnura elegans* (Price et al. 2022); the American grasshopper (Insecta: Orthoptera), *Schistocerca americana* (Childers et al. 2021); the pea aphid (Insecta: Hemiptera), *Acyrthosiphon pisum* (Li et al. 2019); the Australian sheep blowfly (Insecta: Diptera), *Lucilia cuprina* (https://www.ncbi.nlm.nih.gov/assembly/GCF_022045245.1); the silkworm (Insecta: Lepidoptera), *Bombyx mori* (https://www.ncbi.nlm.nih.gov/assembly/GCF_014905235.1); the seven-spotted ladybird (Insecta: Coleoptera), *Coccinella septempunctata* (Crowley et al. 2021b); and the common green lacewing (Insecta: Neuroptera), *Chrysoperla carnea* (Crowley et al. 2021a). Details of the downloaded chromosome-level assemblies are available in Table S1. Furthermore, we downloaded X and autosome assignments of previously published transcripts for the Cristina’s Timema (Insecta: Phasmatodea), *Timema cristinae* (Parker et al. 2022).

Additionally, we downloaded chromosome-level assemblies of the common yellow sally (Insecta: Plecoptera), *Isoperla grammatica* (McSwan et al. 2023), and two springtails (Entognatha: Collembola), *Sinella curviseta* (Zhang et al. 2019) and *Allacma fusca* (Jaron et al. 2022), all of which lack accompanying gene annotations. We therefore downloaded RNA-sequencing reads from the SRA (Sequence Read Archive) on NCBI (Table S1) in order to construct gene annotations for these species. Reads were first trimmed using Trimmomatic v0.39 (Bolger et al. 2014). Reads were then aligned with HISAT2 (Kim et al. 2019), and resulting samfiles were converted to sorted bamfiles using Samtools v.1.16 (Li et al. 2009). GTF files were constructed by StringTie v.2.2.1 (Pertea et al. 2015). Coding sequences were extracted and longest ORFs were identified using Transdecoder v.5.5.0 (Haas, BJ. https://github.com/TransDecoder/TransDecoder). We then extracted the longest isoforms using a custom perl script.

While two of these species, *Allacma fusca* and *Isoperla grammatica*, had identified X chromosomes in their assembly, *Sinella curviseta* did not. To identify the X chromosome in this species, we downloaded paired-end Illumina WGS reads from the SRA from a pool of individuals including both males and females. Reads were aligned using default parameters in Bowtie2 v.2.4.4 (Langmead and Salzberg 2012), and uniquely mapping reads were extracted. We then used soap.coverage (version 2.7.7, https://github.com/gigascience/bgi-soap2/tree/master/tools/soap.coverage/2.7.7) to estimate coverage. The scaffold CM023202.2 had significantly lower coverage than the other five, consistent with the prediction for the X chromosome in a mixed-sex pooled sample (Figure S1). We therefore refer to scaffold CM023202.2 as the X chromosome throughout the manuscript, and rename the other scaffolds as listed in Table S2.

### Detecting orthology to the blue-tailed damselfly and the slender springtail

We selected the blue-tailed damselfly, *I. elegans*, as our reference species because it is the sister lineage to all other insects considered. We therefore refer to *I. elegans* as our reference species, and all other species as our focal species in analyses. However, we also performed all the analyses using the springtail *S. curviseta* as reference, and our result remains largely unchanged.

As a first step, for the seven species with published gene annotations, we processed the translated.cds files and gff files from NCBI (Table S1) through the R package GENESPACE (Lovell et al. 2022), which produces simplified gff files and peptide sequences. For the remaining species with chromosome assemblies but no gene annotation— *A. fusca, S. curviseta*, and *I. grammatica*—we used the longest isoform peptide sequence and modified gtf output from Stringtie, and processed these through GENESPACE as well. For *T. cristinae*, we processed the gff file using a custom python script, and then selected the longest isoform using a custom perl script.

We then used a reciprocal blast approach to define 1-to-1 orthologs between each of our focal species and *I. elegans*, using the processed peptide files from GENESPACE as input. The number of 1:1 orthologs (reciprocal best hits) for comparisons with both *I. elegans* as a reference and *S. curviseta* as a reference are listed in Table 2 and Table S3, respectively. We then counted the number of genes observed as on the X (or Z) in both the focal species and the reference species. To compute the expected number of shared genes for the X chromosome in the reference and each chromosome, we first computed the proportion of 1:1 orthologs on each chromosome in the focal species. This proportion was then multiplied by the number of X-linked 1:1 orthologs in *I. elegans*. In comparisons that use *I. elegans* as the reference, a chromosome was considered homologous if it had a statistically significant excess of shared genes of at least two-fold. As springtails are more distantly related to other taxa, we modified this cutoff to statistical significance and an excess of at least 1.5-fold when using *S. curviseta* as a reference species. Statistical significance was computed using a Monte Carlo approach. Specifically, we randomized chromosome assignments in each comparison for each 1:1 ortholog 100,000 times. The resulting p-values are the proportion of randomizations in which the number of shared orthologs in the randomized dataset are greater or equal to the observed number of shared X-linked (Z-linked) genes. Where p<10^−5^, all simulated datasets have fewer shared orthologs than the observed data.

**Table 2.** Excess of shared gene content between *I. elegans* and the X/Z of the focal species.

| Order | Focal Species | 1:1<br>Orthologs | Chr | Expected<br>Shared X | Observed<br>Shared X | Exp/<br>Obs | p-<br>value |
| --- | --- | --- | --- | --- | --- | --- | --- |
| Plectoptera | *I. grammatica* | 3458 | X<sub>1</sub> | 9 | 56 | 6.2 | <10<sup>-5</sup> |
| Plectoptera | *I. grammatica* | 3458 | X<sub>2</sub> | 10 | 27 | 2.7 | <10<sup>-5</sup> |
| Orthoptera | *S. americana* | 5995 | X | 47 | 186 | 4.0 | <10<sup>-5</sup> |
| Phasmatodea | *T. cristinae* | 2166 | X | 7 | 33 | 4.7 | <10<sup>-5</sup> |
| Lepidoptera | *B. mori* | 4625 | Z | 13 | 3 | 0.2 | 0.9999 |
| Diptera | *L. cuprina* | 4076 | X | 1 | 7 | 7.0 | 3x10<sup>-5</sup> |
| Coleoptera | *C. septempunctata* | 4904 | X | 23 | 100 | 4.3 | <10<sup>-5</sup> |
| Neuroptera | *C. carnea* | 5147 | X | 23 | 94 | 4.1 | <10<sup>-5</sup> |
| Hemiptera | *A. pisum* | 4320 | X | 33 | 86 | 2.6 | <10<sup>-5</sup> |
| Collembola | *S. curviseta* | 3570 | X | 46 | 116 | 2.5 | <10<sup>-5</sup> |
| Collembola | *A. fusca* | 3318 | X<sub>1</sub> | 56 | 109 | 1.9 | <10<sup>-5</sup> |
| Collembola | *A. fusca* | 3318 | X<sub>2</sub> | 36 | 35 | 1.0 | 0.624 |

### Detecting fusions and fissions to sex chromosomes across the insect phylogeny

Fusions and fissions are common in sex-chromosome evolution (Bachtrog et al. 2014; Pennell et al. 2015), and may have played an important role in insects. In order to try to detect these events across the phylogeny, we then identified the regions of the genome in our reference species (*I. elegans* and *S. curviseta*) which were homologous to the X chromosome of the focal species. This is simply the reciprocal analyses described above. We again defined excess as two-fold using *I. elegans*, and 1.5-fold using *S. curviseta*, and computed significance using the Monte Carlo approach described above.

### Synteny detection in insect genomes

Finally, we selected four insect species to examine synteny across the phylogeny: *S. americana, C. septempunctata, C. carnea*, and *I. elegans*. The synteny map was constructed using GENESPACE v0.94 (Lovell et al. 2022).

## Results and Discussion

### The X chromosome is conserved across insects

We first systematically tested whether the same sex chromosome was used throughout insects. To do so, we compared the gene content of the X of clades for which homology to the dipteran X had been detected in previous studies (Odonata, Coleoptera, Orthoptera, Hemiptera, and Diptera itself; Chauhan et al. 2021; Xinghua et al. 2022) as well as four additional orders (Phasmatodea, Neuroptera, Plecoptera, and the ZW Lepidoptera). All but one genome assembly were chromosome-level and had annotated sex chromosomes. For the Phasmatodean *T. cristinae*, only scaffolds were available, but these were previously assigned as X-linked or autosomal based on their male and female genomic coverage (Parker et al. 2022). We used the damselfly *I. elegans* (Odonata) as the reference species, as it is sister to all other insect lineages considered.

In all seven insect orders with heterogametic XY or XO sex chromosomes, we detected homology with the X chromosome of the damselfly, *I. elegans* (p<0.0001 in every case, Monte Carlo randomization, Figure 1; Table 2; Figure S2). Further, in the order Plecoptera, *I. grammatica* has two X chromosomes (X_1_ and X_2_), both of which are homologous to the *I. elegans* X chromosome. Importantly, the Lepidopteran Z chromosome shows no homology to the *I. elegans* X chromosome (Figure 1; Table 2; Figure S2), which is consistent with previous results (Chauhan et al. 2021), and suggests that the switch in heterogametey that occurred in this lineage was associated with a turnover in sex chromosome. Finally, in *I. grammatica, S. americana*, C. *septempunctata*, and *C. carnea*, an autosome was also homologous to the X chromosome of *I. elegans*, though in all cases it had a smaller excess of shared gene content than the X chromosome (Figure 1; Table S4).

**Figure 1.**
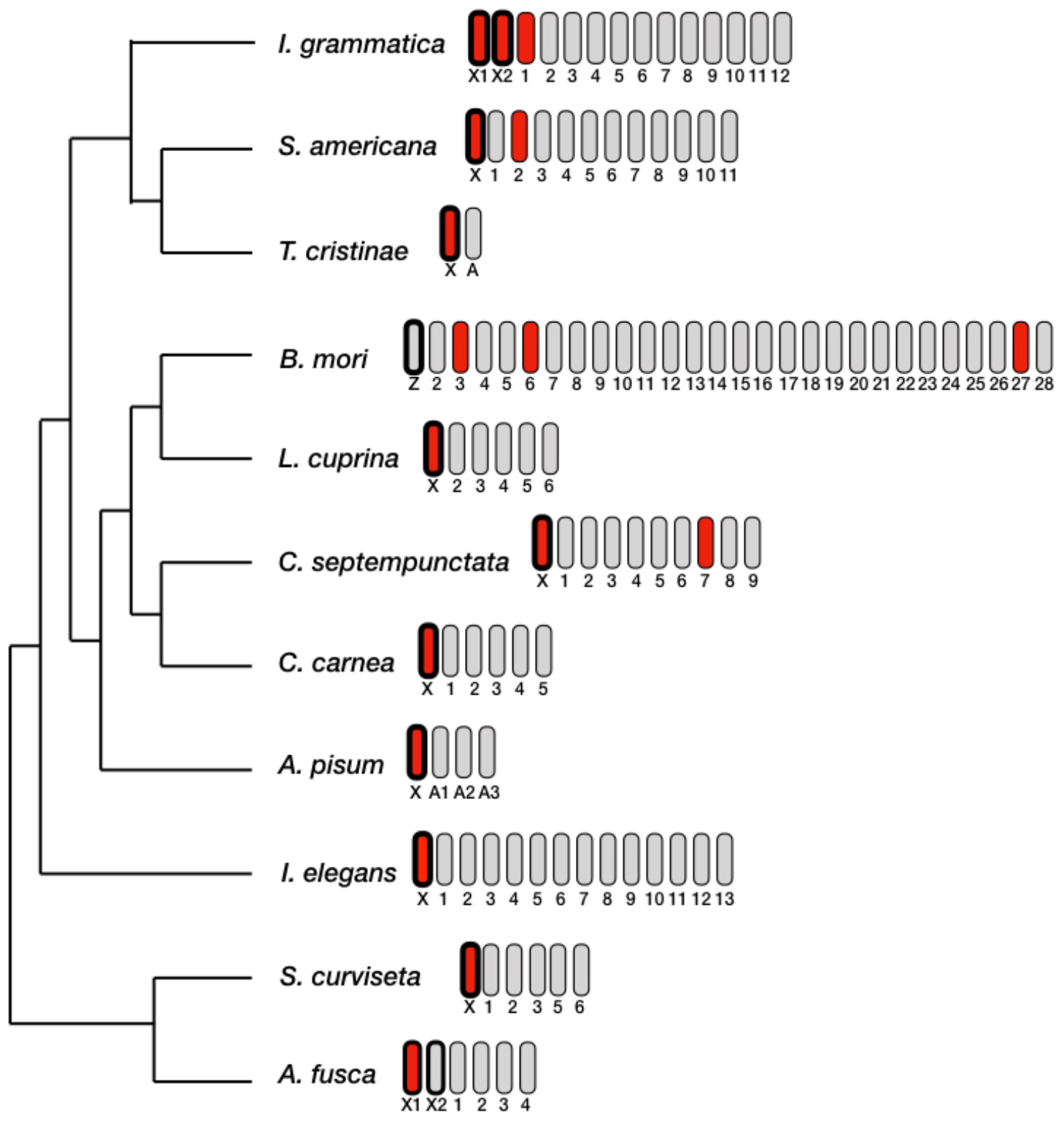
Homology of X chromosomes across insects. Phylogeny is adapted from Blackmon et al (2017). Idiograms for each species are shown, with chromosomes homologous to the *I. elegans* X chromosome colored red, and those not homologous colored grey. Sex chromosomes for each species are outlined in black.

We also implemented a synteny analysis using GENESPACE. We were able to detect conservation of the X chromosome across the phylogeny when using four species representing distant insect lineages: *I. elegans, S. americana, C. carnea*, and *C septempunctata* (Figure S4). However, the signal disappears when all species are used, as few syntenic blocks are shared by all species.

### The ancestral insect X chromosome is shared with springtails (Entognatha: Collembola)

We then investigated whether the origin of the X may have predated the appearance of insects altogether. To do so, we compared the *I. elegans* genome to the closest outgroup of insects, the springtails (Collembola). We selected two species with genome assemblies, *Sinella curviseta* and *Allacma fusca*, which are at least 193 million years diverged from each other (Kumar et al. 2017; Yu et al. 2021). We then downloaded RNA-seq reads for both species, as neither genome had gene annotations, and constructed transcriptome assemblies to use in downstream analyses. The *A. fusca* annotation has two X chromosomes, X_1_ and X_2_ (Jaron et al. 2022). In contrast, no information is available as to what chromosome is the X in *S. curviseta*. We therefore downloaded pooled male and female genomic DNA from the SRA, and performed a coverage analysis to identify the X chromosome. We detected homology between the X chromosome of *I. elegans* and the inferred X of *S. curviseta*, as well as the X_1_ chromosome of *A. fusca* (Table 1; Figure 1), supporting a more ancient origin of the X than of insects.

### Potential fusion of *I. elegans* chromosome 2 to the X in distant lineages

It has been suggested that it may be selectively beneficial to recruit specific chromosomes or genomic regions as sex chromosomes, if these regions harbor genes involved in sex-determination or a pre-existing excess of sexually antagonistic gene content (Marshall Graves and Peichel 2010; Jeffries et al. 2018; Toups et al. 2019). To investigate this, we performed the reciprocal homology analysis to that described above, i.e. we asked which autosomes of *I. elegans* became X-linked in each of the other lineages. We then tested if one or more autosomes were recruited onto the X more often than expected by chance. We find that chromosome 2 in *I. elegans* is sex-linked in the phylogenetically distantly related taxa of *I. grammatica, T. cristinae, L. cuprina*, and *C. carnea* (Table S4). If chromosome 2 became sex-linked independently through fusion events in these lineages, then it did more than expected by chance within insects (binomial test, p<0.05). However, if we extend our analysis to include Collembola, it is no longer statistically significant (binomial test, p=0.07282). While it is tempting to speculate that the sex-linkage of (a part of) chromosome 2 may be advantageous, it is also possible the region of the genome corresponding to *I. elegans* chromosome 2 was incorporated into the X chromosome of insects and subsequently lost in *C. septempunctata, A. pisum*, and *S. americana*. Finally, rearrangements such as fissions and fusions are common in Coleoptera (Bracewell et al. 2023) and Hemiptera (Mathers et al. 2021), and the above result may simply occur because of the taxa chosen within these orders. More extensive sampling would be needed to distinguish between these three hypotheses. Interestingly, the X chromosome of the slender springtail, *S. curviseta*, and the X_1_ chromosome of the globular springtail, *A. fusca*, is homologous to chromosome 7 in *I. elegans*, in addition to the X chromosome. While we cannot in this case distinguish between a fusion of the X and autosome 7 in Collembola or a loss of part of the ancestral hexapod X (now corresponding to chromosome 7) in the common ancestor of insects, this further emphasizes the long-term conservation of gene content within Collembola, and arthropods in general.

### Few genes shared among all Xs

Figure 1 suggests that while the ancestral X chromosome has remained sex-linked throughout insect history, substantial chromosomal reshuffling has occurred, as multiple chromosomes correspond to the *I. elegans* X in the other species. Even if the same chromosome is used as the X in all insect orders, extensive gene movement within and between chromosomes, combined with rare translocations, could lead to different gene sets remaining X-linked throughout the insect phylogeny. On the other hand, selection may have preserved a substantial set of genes with sex-specific roles on the X. To investigate this, we compared the gene content of the X of the different orders. We identified a total of 2253 1:1 orthologs between the Plecoptera, Orthoptera, Diptera, Coleoptera, Neuroptera, Hemiptera, Odonata, and Collembola. Of these, only 1 was shared across the entire phylogeny on the X chromosome (Figure 2). This is largely because the X chromosome in Diptera is considerably smaller than the other orders, with only nine 1:1 orthologs present on the X (out of 207 total X genes). When Diptera is removed from the analysis, 14 genes are shared between the X chromosomes of insects and springtails. Importantly, the X chromosome of all orders are homologous with the Collembola X chromosome (Figure S3; Table S3; Table S5). Despite these constraints, it is clear that even when the same chromosome is used as the X over long periods of time, the majority of X-linked genes are not necessarily shared.

**Figure 2.**
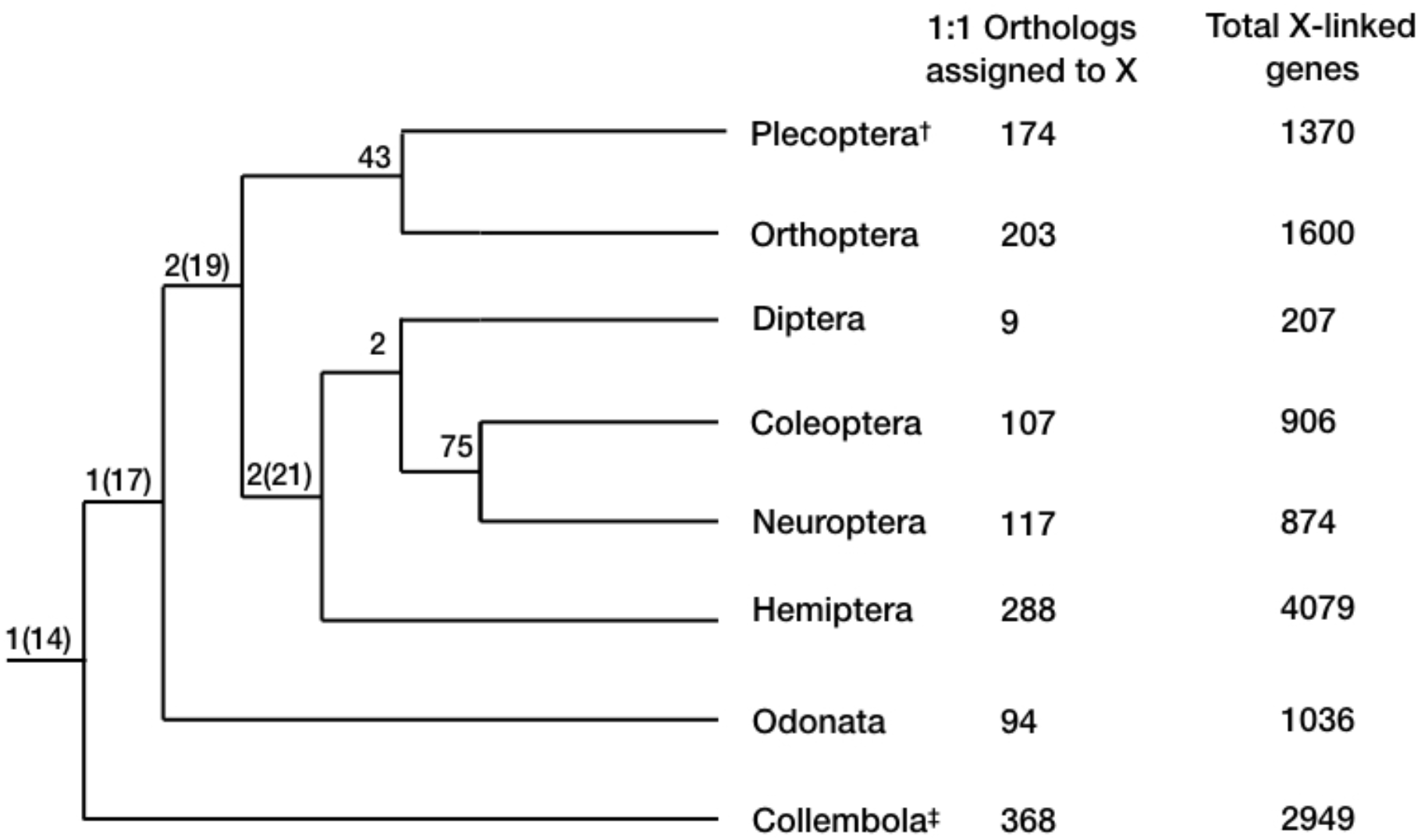
Shared gene content across the insect and springtail phylogeny. Values in parentheses are computed excluding the Dipteran X chromosome. ^†^Values for Plecoptera are combined for X_1_ and X_2_, as both are homologous to the *I. elegans* X chromosome. ‡Values for Collembola only consider *S. curviseta*.

Finally, it is clear from Figure 2 that the small number of genes found on the Diptera X chromosome is a derived feature, which may help explain a paradoxical observation: how does an X chromosome that has been conserved since the origin of insects suddenly start undergoing turnover repeatedly in flies and mosquitoes? Such turnover is thought to be prevented by the damaging consequences of putting a highly specialized sex chromosome in an autosomal context (for instance, this could lead to dosage compensation being active despite no longer being needed). The resulting fitness effect is likely proportional to the number of genes affected, and the reduction in gene content of the dipteran X may have been a pre-requisite for its reversal to an autosome.

## Conclusions

We present evidence that the X chromosome is shared among at least eight insect orders and the outgroup Collembola, and originated prior to the evolution of Class Insecta itself. This is an addition to the German cockroach, *Blatella germanica* (Insecta: Blattodea), whose X is homologous to element F in Dipterans (Meisel et al. 2019), which we omitted from this analysis because there are currently no chromosome-level genome assemblies. This suggests that the insect X chromosome is at least 459 MYA (CI: 425.4 −478.1), which is the most ancient sex chromosome identified to date. Further, we suggest that the shrinking X chromosome content of the Dipteran X chromosome allowed it to escape the “evolutionary trap” of highly differentiated sex chromosomes, which led to high rates of sex-chromosome turnover among fly lineages (Vicoso and Bachtrog 2015).

