## Supplementary Figures and Tables for "The X chromosome of insects predates the origin of Class Insecta"

**Supplementary Material**

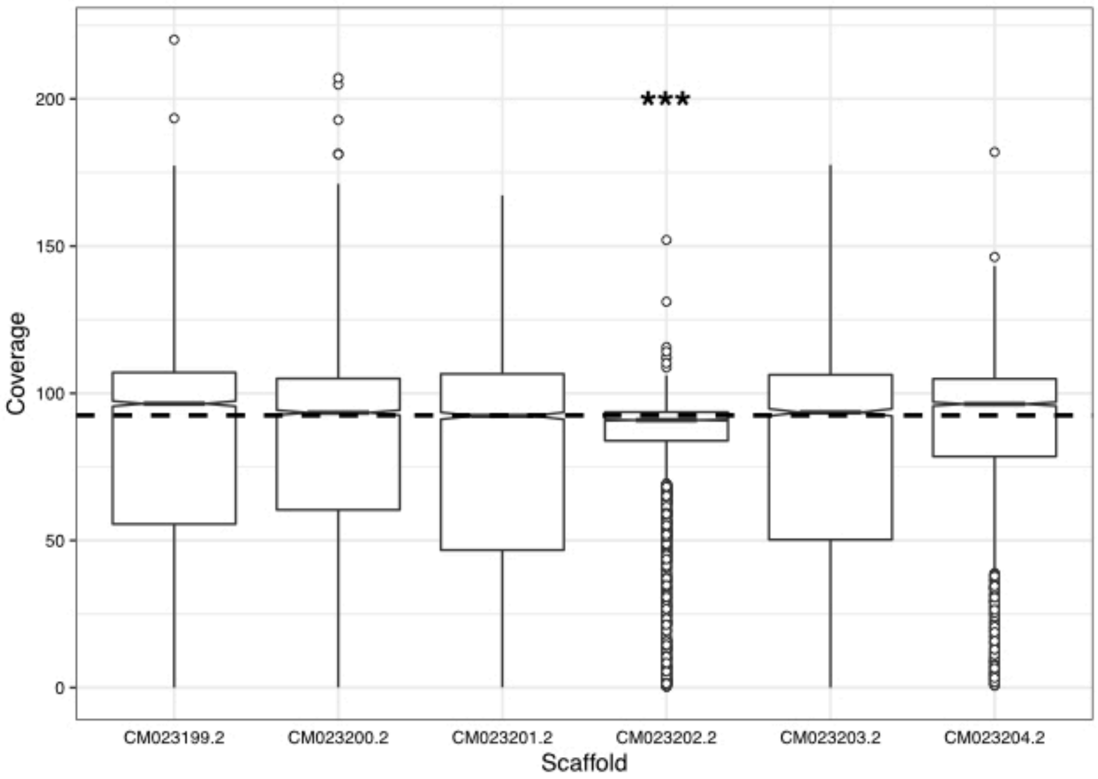


**Figure S1.** Coverage of a pool mixed-sex individuals of *S. curviseta*. Scaffold CM023202.2 has significantly lower coverage than all other chromosomes, and a coverage lower than the genome median (horizontal dashed line). We therefore assigned it as the X chromosome in our downstream analyses.

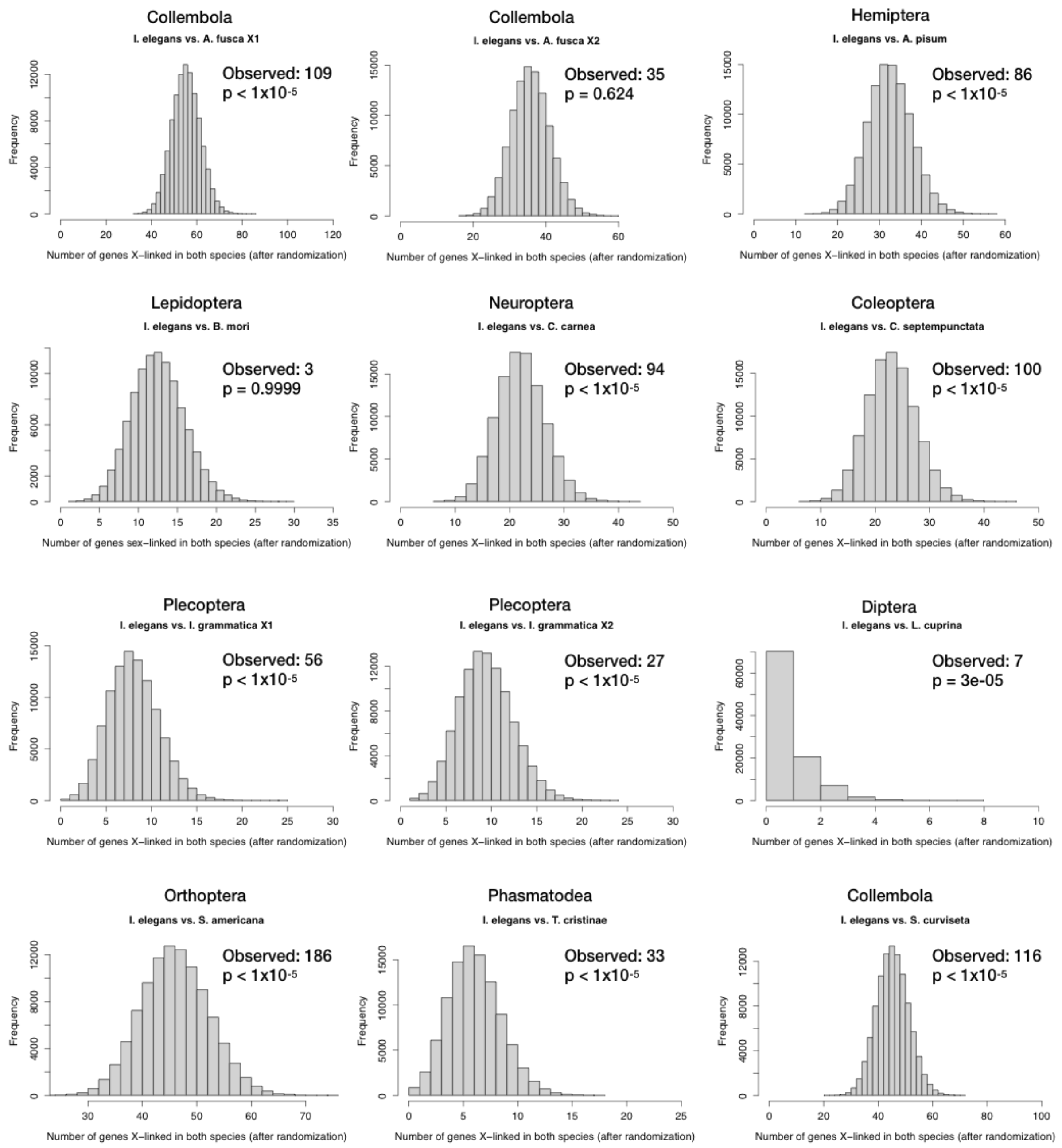


**Figure S2**. Significance tests for excess shared gene content between the X chromosome of the blue-tailed damselfly, *I. elegans*, and focal species across 8 insect orders and the outgroup Collembola. Tests were performed for both $X_1$ and $X_2$ chromosomes in the stonefly, *I. grammatica*, and the springtail, *A. fusca*.

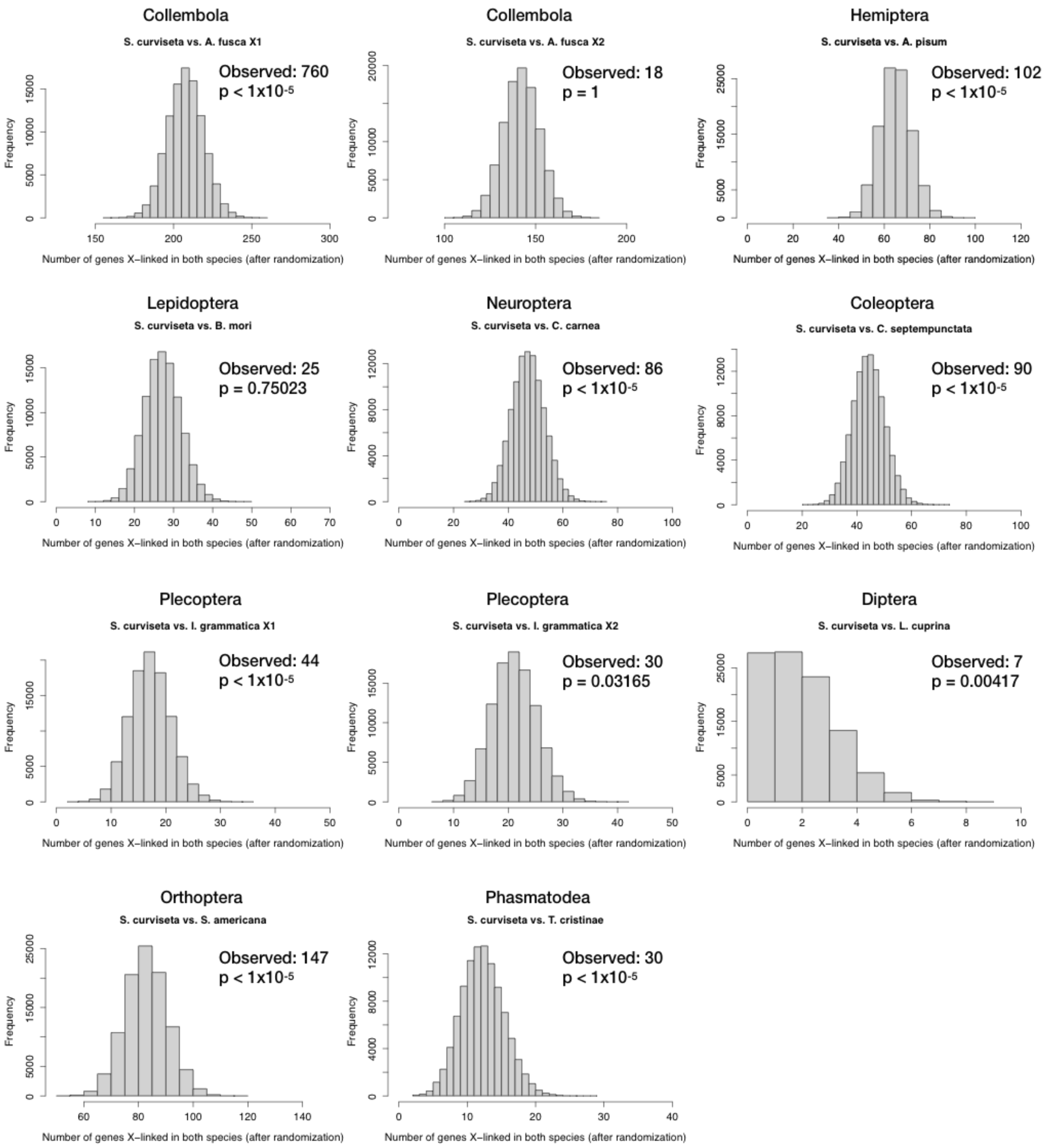


**Figure S3**. Significance tests for excess shared gene content between the X chromosome of the slender springtail, *S. curviseta*, and focal species across 8 insect orders and the outgroup Collembola. Tests were performed for both $X_1$ and $X_2$ chromosomes in the stonefly, *I. grammatica*, and the springtail, *A. fusca*.

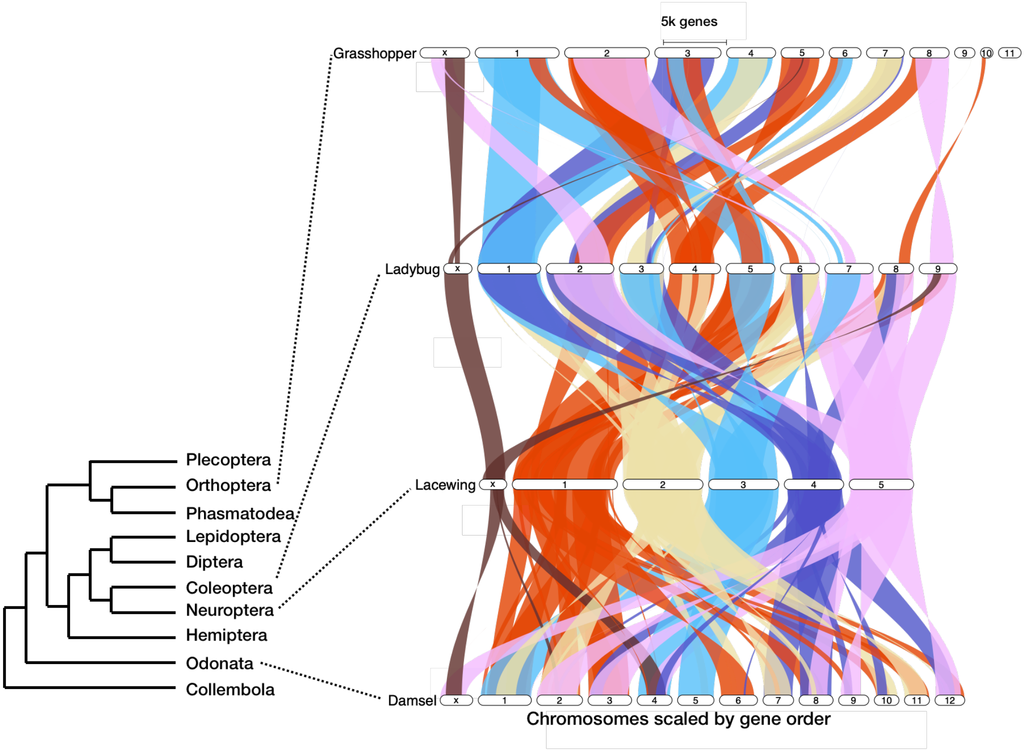


**Figure S4.** Synteny is conserved on the X chromosome across class Insecta. Phylogeny is adapted from Blackmon et al. (2017). Synteny is plotted with respect to *C. carnea* (lacewing; Neuroptera) using GENESPACE (Lovell et al. 2022).

**Table S1.** Genome assemblies and RNA-seq and DNA-seq included in the analyses.

| Species | Order | genome version | RNA-seq | DNA-Seq |
|---|---|---|---|---|
| *Isoperla grammatica* | Plecoptera | GCA_945910005.1 | ERR9881692 | NA |
| *Schistocerca americana* | Orthoptera | GCF_021461395.2 | NA | NA |
| *Bombyx mori* | Lepidoptera | GCF_014905235.1 | NA | NA |
| *Lucilia cuprina* | Diptera | GCF_022045245.1 | NA | NA |
| *Coccinella septempunctata* | Coleoptera | GCF_907165205.1 | NA | NA |
| *Chrysoperla carnea* | Neuroptera | GCF_905475395.1 | NA | NA |
| *Acyrthosiphon pisum* | Hemiptera | GCF_005508785.2 | NA | NA |
| *Sinella curviseta* | Collembola | GCA_004115045.3 | SRR7948082 | SRR7948081 |
| *Allacma fusca* | Collembola | GCA_947179485.1 | SRR1653171 | NA |
| *Ischnura elegans* | Odonata | GCF_921293095.1 | NA | NA |

**Table S2.** Scaffold ID in *S. curviseta* genome assembly, and its corresponding ID in analyses.

| Scaffold ID | Chromosome ID |
|---|---|
| CM023199.2 | 1 |
| CM023200.2 | 2 |
| CM023201.2 | 3 |
| CM023202.2 | X |
| CM023203.2 | 5 |
| CM023204.2 | 6 |

**Table S3.** Excess of shared gene content between *S.curviseta* and the X/Z of the focal species.

| **Order** | **Focal Species** | **1:1 Orthologs** | **Chr** | **Expected Shared X** | **Observed Shared X** | **Exp/ Obs** | **p-value** |
|---|---|---|---|---|---|---|---|
| Plectoptera | *I. grammatica* | 2739 | $X_1$ | 18 | 44 | 2.4 | $<10^{-5}$ |
| Plectoptera | *I. grammatica* | 2739 | $X_2$ | 22 | 30 | 1.4 | 0.0317 |
| Orthoptera | *S. americana* | 3765 | X | 83 | 147 | 1.8 | $<10^{-5}$ |
| Phasmatodea | *T. cristinae* | 1484 | X | 13 | 30 | 2.3 | $<10^{-5}$ |
| Lepidoptera | *B. mori* | 3280 | Z | 28 | 25 | 0.9 | 0.7502 |
| Diptera | *L. cuprina* | 2945 | X | 2 | 7 | 3.5 | 0.0042 |
| Coleoptera | *C. septempunctata* | 3378 | X | 45 | 90 | 2.0 | $<10^{-5}$ |
| Neuroptera | *C. carnea* | 3340 | X | 48 | 86 | 1.8 | $<10^{-5}$ |
| Hemiptera | *A. pisum* | 3040 | X | 65 | 102 | 1.6 | $<10^{-5}$ |
| Collembola | *A. fusca* | 5021 | $X_1$ | 208 | 760 | 3.7 | $<10^{-5}$ |
| Collembola | *A. fusca* | 5021 | $X_2$ | 143 | 18 | 0.1 | 1 |

**Table S4.** Orthology relationships between the X chromosomes of a representative species of eight insect species and two Collembola species with the blue-tailed damselfly, *I. elegans*. Chromosomes are ordered according to their proportion of shared homology to the X chromosome, with the highest proportion first. §A full analysis is not possible with T. cristinae, as we lack a genome assembly.

| **Order** | **Species** | **Chromosomes with an excess of homologs of *I. elegans* X-linked genes** | **X (Z) homolog in *I. elegans*** |
|---|---|---|---|
| Plecoptera | *I. grammatica* | $X_1$,$X_2$,1 | $X_1$: X,7<br>$X_2$: 4,X,2 |
| Orthoptera | *S. americana* | X,2 | X,9,12 |
| Phasmatodea | *T. cristinae* | X§ | X,2 |
| Lepidoptera | *B. mori* | 3,27,6 | 4,8 |
| Diptera | *L. cuprina* | X | X,2,10 |
| Coleoptera | *C. septempunctata* | X,7 | X |
| Neuroptera | *C. carnea* | X,5 | X,4,2 |
| Hemiptera | *A. pisum* | X | 11,X |
| Collembola | *S. curviseta* | X | 7,X |
| Collembola | *A. fusca* | $X_1$ | $X_1$: 7,X,3.5<br>$X_2$: none |

**Table S5.** Orthology relationships between the X chromosomes of a representative species of 8 insect species and Collembola species with the slender springtail, *S. curviseta*. Chromosomes are ordered according to their proportion of shared homology to the X chromosome, with the highest proportion first. §A full analysis is not possible with *T. cristinae*, as we lack a genome assembly.

| **Order** | **Species** | **Chromosomes with an excess of homologs of *S. curviseta* X-linked genes** | **X (Z) homolog in *S. curviseta*** |
|---|---|---|---|
| Plecoptera | *I. grammatica* | $X_1$,4,11,3 | $X_1$: X<br>$X_2$: none |
| Orthoptera | *S. americana* | 8,X,7 | X,2 |
| Phasmatodea | *T. cristinae* | X§ | X |
| Lepidoptera | *B. mori* | 16,14,13,26,<br>27,23 | None |
| Diptera | *L. cuprina* | X | X |
| Coleoptera | *C. septempunctata* | X,9,3 | X |
| Neuroptera | *C. carnea* | X | X |
| Hemiptera | *A. pisum* | X | 5,X |
| Collembola | *A. fusca* | $X_1$ | $X_1$: X<br>$X_2$: 3 |
